# Vimentin intermediate filaments support autophagosome biogenesis at ER-endosomes contact sites in response to starvation

**DOI:** 10.64898/2025.12.02.691851

**Authors:** Quentin Frenger, Damiana Lecoeuche, Cédric Delevoye, Etienne Morel

**Author notes:** Lead contact, corresponding Author: Etienne Morel.

## Abstract

Macroautophagy (autophagy) is a fundamental catabolic process requiring the biogenesis of the autophagosome to support cell survival during stress. While the roles of F-actin and microtubule cytoskeleton in autophagy are well established, the contribution of intermediate filaments (IFs) remains poorly understood. Here, we investigated the role of the type III IF vimentin in supporting the early steps of starvation-induced autophagy. We demonstrate that starvation triggers a rapid, perinuclear compaction of vimentin IFs, correlating with transient phosphorylation at serine 56 and enhanced overlap with the endoplasmic reticulum (ER). We reveal that autophagic proteins accumulate at the vimentin/ER interface, physically connecting the autophagosome biogenesis machinery to the vimentin IF network. Knock-out or pharmacological perturbation of vimentin-IFs dynamics using Withaferin-A significantly impairs starvation-induced autophagic flux. Mechanistically, we reveal that vimentin IFs are essential coordinators for the mobilization of endosome-ER-membrane contact sites (EERCS), a critical hub for autophagosome nucleation. Together, our findings uncover a novel role for vimentin IFs as a dynamic cytoskeletal coordinator that spatially organizes membrane contact sites to promote the efficient initiation of autophagosome biogenesis in response to nutrient stress.

**Summary statement:** This study reveals that vimentin intermediate filaments rapidly reorganize to mobilize ER-endosome contact sites, establishing a critical spatial platform for starvation-induced autophagy initiation.

## Introduction

During stress response, organelles are dynamically mobilized, shaped, and repositioned to support cellular homeostasis (Da Graça et al., 2024). In this context, macroautophagy (hereinafter autophagy) is a major catabolic process engaged to support cell survival in response to nutrient starvation. Autophagy is coordinated by autophagy-related (ATG) protein activity and is dysregulated in several diseases, highlighting the importance of a precise understanding of its regulation (Klionsky et al., 2021; Mizushima & Levine, 2020).

Under nutrient deprivation, autophagy is initiated by the ULK complex (FIP200, ATG13, ATG101, ULK1/2) at the Endoplasmic Reticulum (ER) membrane. This complex activates the PI3KC3-C1 complex (VPS34, VPS15, Beclin-1, ATG14, NRBF2) through ULK1-dependent phosphorylation (Kim et al., 2011; Russell et al., 2013). The VPS34 kinase activity generates phosphatidylinositol-3-phosphate (PI3P) at the surface of the ER in a peculiarly curved zone called the omegasome. Then, PI3P-binding proteins, such as WIPI family β-propeller proteins (e.g. WIPI2) and zinc-finger FYVE domain-containing protein 1 (DFCP1), are recruited to this specific area to support the mobilization of ATG proteins for the assembly of the pre-autophagic structure, termed the phagophore (Axe et al., 2008; Polson et al., 2010). Phagophore formation begins at ER–membrane contact-sites, including ER-mitochondria, ER-plasma membrane and ER-endosomes (Da Graça et al., 2025; Hamasaki et al., 2013; Nascimbeni et al., 2017). Here, several ATGs are recruited to initiate and sustain the *de novo* biogenesis of the phagophore. Importantly, the ATG12–ATG5-ATG16L1 complex is recruited via WIPI2 to catalyze the final step of the conjugation of ATG8 proteins (including LC3) to phosphatidylethanolamine, also called membrane ATG8ylation, a hallmark of autophagosome biogenesis (Deretic et al., 2024). Lipidated LC3 then interacts with receptors, such as p62, to ensure autophagosome closure around its cargo (Dooley et al., 2014; Pankiv et al., 2007; Young et al., 2006). Following closure, the autophagosome is trafficked to lysosomes for fusion and content degradation, enabling component recycling.

The intricate steps of autophagosome biogenesis rely on the precise spatial and temporal coordination of membrane dynamics. This coordination is largely governed by the mammalian cytoskeleton, which comprises F-actin, microtubules, and IFs. While the roles of F-actin and microtubules in organelle positioning and transport are well-established (Barlan & Gelfand, 2017; Brown, 1999), the specific contribution of IFs remains less understood. IFs components are ubiquitously expressed and can form mesh-like structures that support cellular architecture and mechanically position organelles (Eriksson et al., 2009; Guo et al., 2025; Pasolli et al., 2025). For instance, our lab recently demonstrated that keratin IFs in skin keratinocytes form structural cages around lysosome-like pigment organelles, maintaining their perinuclear distribution to protect the genome from UV damage (Benito-Martínez et al., 2025). Similarly, type III IFs composed of vimentin have been shown to encage pathogens such as *C. trachomatis* or the Dengue virus (Kumar & Valdivia, 2008; Teo & Chu, 2014). Importantly, vimentin IFs are also tightly associated with the ER and regulate the subcellular localization of the nucleus, aggresomes, mitochondria, lipid droplets, and lysosomes (Morrow et al., 2020; van Bodegraven et al., 2025; Biskou et al., 2019; Cremer et al., 2023; Pasolli et al., 2025).

Our recent work identified that endosome-ER membrane contact-sites (EERCS) as a crucial platform for *de novo* phagophore biogenesis in response to starvation (Da Graça et al., 2025). Given that vimentin IFs are regulators of organelle positioning, we investigated their role in this sequence. Here, we demonstrate that starvation triggers a rapid (within 15 minutes) compaction of the vimentin IFs in the perinuclear area, driving its accumulation at the ER interface. We show that this rapid transient reorganization is partly controlled by a temporally regulated phosphorylation of vimentin at serine 56. Importantly, this reorganized vimentin assembly concentrates key pre-autophagic markers (DFCP1, LC3B, ATG13) at specific ER domains. We show that impairing vimentin dynamics, either pharmacologically (Withaferin-A) or genetically (KO), abolishes the rapid mobilization of EERCS and blocks autophagic flux. Collectively, our findings unveil vimentin as a dynamic coordinator that spatially orchestrates membrane contact sites to nucleate autophagosomes.

## Results

### Vimentin network is compacted in the perinuclear region in response to starvation

The cellular response to nutrient stress requires a precise spatiotemporal mobilization of organelles. Given the established role of intermediate filaments in regulating cellular spatialization and organelle positioning, we hypothesized that the vimentin IFs would undergo spatial modulation to support starvation-induced autophagy. To test this, we starved HeLa cells using Earl’s Balanced Salt Solution (EBSS) for 15 or 60 minutes and analyzed vimentin organization by STED super-resolution microscopy following immunofluorescence staining with an anti-vimentin antibody. As shown in Figure 1A, in control (untreated) cells, the vimentin IF network was broadly distributed throughout the cytoplasm, displaying its typical extended filamentous pattern (Battaglia et al., 2018). In contrast, upon starvation, vimentin IF accumulated predominantly in the perinuclear region, as indicated by a marked decrease in signal intensity in the cell periphery (Fig. 1A, 15 min and 60 min starvation). To quantify this redistribution, we processed spinning disk microscopy acquisitions and generated segmentation masks to define the filament network morphology in cells divided in two equal bins corresponding to the peripheral and perinuclear area (Fig. 1B). Quantification revealed a significant increase in vimentin signal within the perinuclear zone after 15 or 60 min. of starvation compared with untreated cells (Fig. 1C, D). We further analyzed network morphology using the CellProfiler compactness metric. In this analysis, high values reflect a highly complex, spatially dispersed filament network, whereas a reduction in this index indicates a transition toward a denser shape. Consistently, the compactness value significantly decreased upon starvation (Fig. 1E, F), confirming the spatial compaction of the vimentin network into the perinuclear region. Finally, given that starvation triggers extensive ER remodeling, such as the conversion of peripheral tubules into perinuclear sheets (Jang et al., 2022) (Lu et al., 2020), and since vimentin regulates ER morphodynamics, (Cremer et al., 2023; Pasolli et al., 2025), we analyzed the co-distribution of vimentin and ER networks. As shown in Supplementary Figure 1, the vimentin to Sec61β^RFP^ signal (ER membrane) overlap is increased in response to starvation, suggesting that the local vimentin IFs reorganization observed in response to nutrient stress correlates, at least in part, with ER membrane. Together, these data demonstrate that vimentin undergoes rapid and pronounced perinuclear reorganization and ER association upon nutrient starvation.

**Figure 1.**
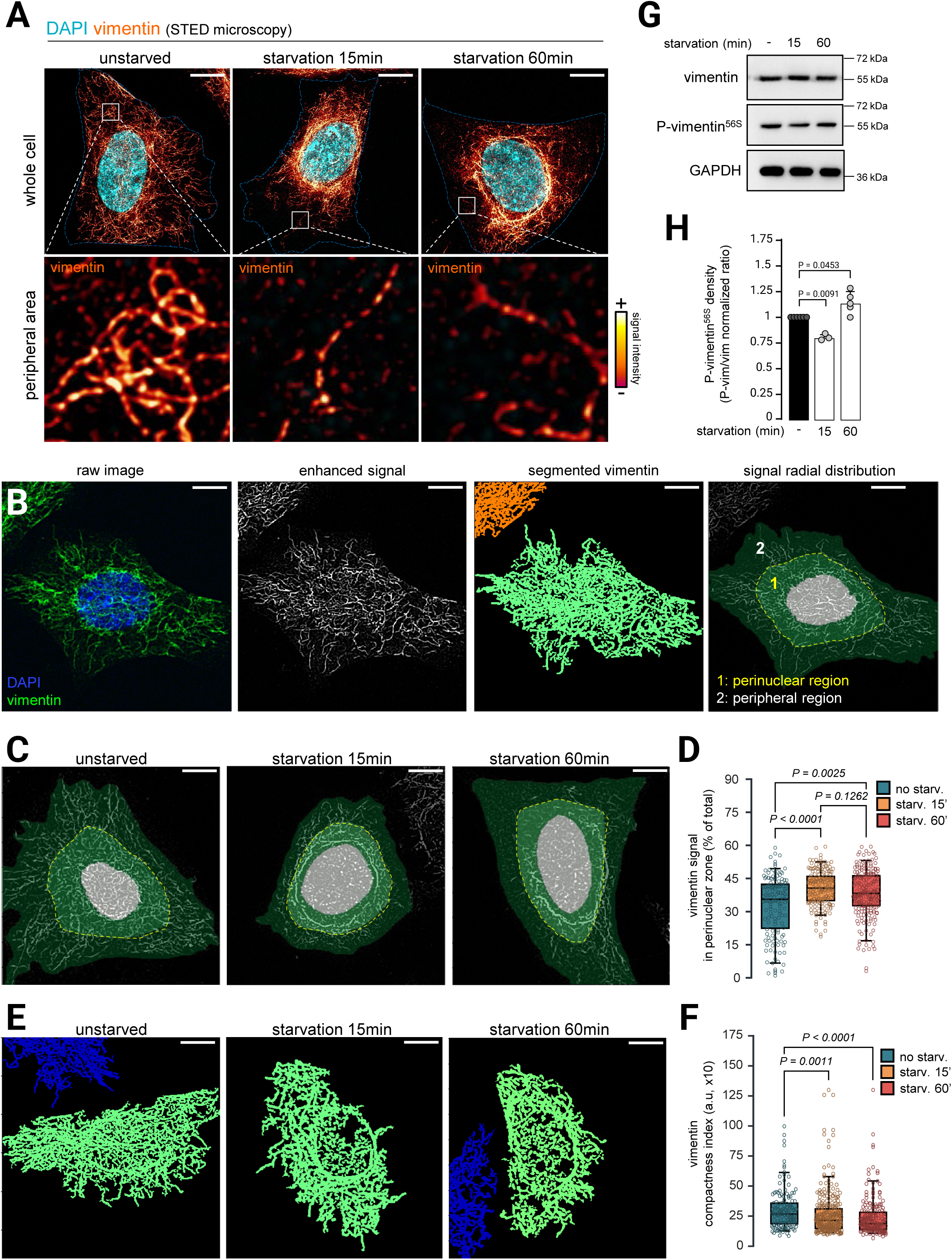
The vimentin network is rearranged in the perinuclear region in response to starvation. HeLa cells were analyzed in basal conditions or after starvation with EBSS for 15 or 60 minutes, unless otherwise noted. (**A**) Stimulated-emission-depletion (STED) images of cells labeled with DAPI and an anti-vimentin antibody. Inserts show the vimentin network in the peripheral area of the cytoplasm. Scale bar = 10 µm. (**B**) Illustrations of the technical workflow employed to segment and quantify the morphology of the vimentin network. From left to right: raw image of cells labeled with DAPI and an anti-vimentin antibody, enhanced vimentin signal obtained after pre-processing using CellProfiler’s “Enhance Or Suppress Features” module, mask of the segmented vimentin network, division of the cell cytoplasm into perinuclear region (1, yellow dots) or the peripheral region (2) obtained with CellProfiler’s “Measure Object Intensity Distribution” module. (**C**) Representative confocal images of the cytoplasmic distribution of the vimentin network of cells labeled with an anti-vimentin antibody. (**D**) Quantification of the proportion of the vimentin signal in the perinuclear region (light green, yellow dots). N=4 independent experiments; n=144 (basal), 186 (15 min), 216 (60 min) cells. *P*-values calculated with Kruskal-Wallis test with Dunn’s multiple comparisons test. (**E**) Representative images of the segmented vimentin network (green) of cells labeled with DAPI and an anti-vimentin antibody. (**F**) Quantification of compactness index (calculated as perimeter²/(4π · area)) of the segmented vimentin network. N=3 independent experiments; n=143 (basal), 298 (15 min), 197 (60 min) cells. *P*-values calculated with Kruskal-Wallis test with Dunn’s multiple comparisons test. (**G**) Representative western blot analysis of vimentin, P-vimentin56S and GAPDH expression of HeLa cells in basal or EBSS-starved (15, 60 or 360 minutes) conditions. (**H**) Quantification of western blot analysis shown in (**G**). P-vimentin56S density was normalized with vimentin density. N=5 (basal), 3 (15 min), 5 (60 min), 5 (360 min) independent experiments. *P*-values calculated with two-tailed Mann-Whitney tests.

### Vimentin phosphorylation at serine 56 is dynamically regulated in response to starvation

To further investigate the mechanisms underlying vimentin reorganization upon nutrient deprivation, we examined the phosphorylation status of vimentin at serine 56, a modification previously shown to regulate vimentin IFs network dynamics (Li et al., 2006). Immunofluorescence analysis revealed that in untreated cells, P-Vimentin^S56^ displayed a punctate cytoplasmic distribution. Strikingly, after only 15 minutes of starvation, this signal became concentrated in perinuclear structures (Fig. S2A, B). Notably, this phenotype was transient, as the signal reverted to a punctate pattern after 60 minutes of starvation (Sup. Fig. 2A, B). To assess global changes in P-Vimentin^S56^ levels, we performed western blot analyses after 15 minutes or 60 minutes of starvation. Total vimentin protein levels remained unchanged across all conditions. In contrast, P-Vimentin^S56^ levels showed dynamic regulation over time: a modest but significant decrease after 15 minutes, followed by a transient increase at 60 minutes (Fig. 1G, H). This temporal pattern likely reflects the transient activation of a vimentin-specific response to starvation. Together, these immunofluorescence and western blot analyses indicate that vimentin Ser56 phosphorylation is dynamically regulated during nutrient stress. This transient phosphorylation correlates with the rapid perinuclear rearrangement of the vimentin network observed upon starvation.

**Figure 2.**
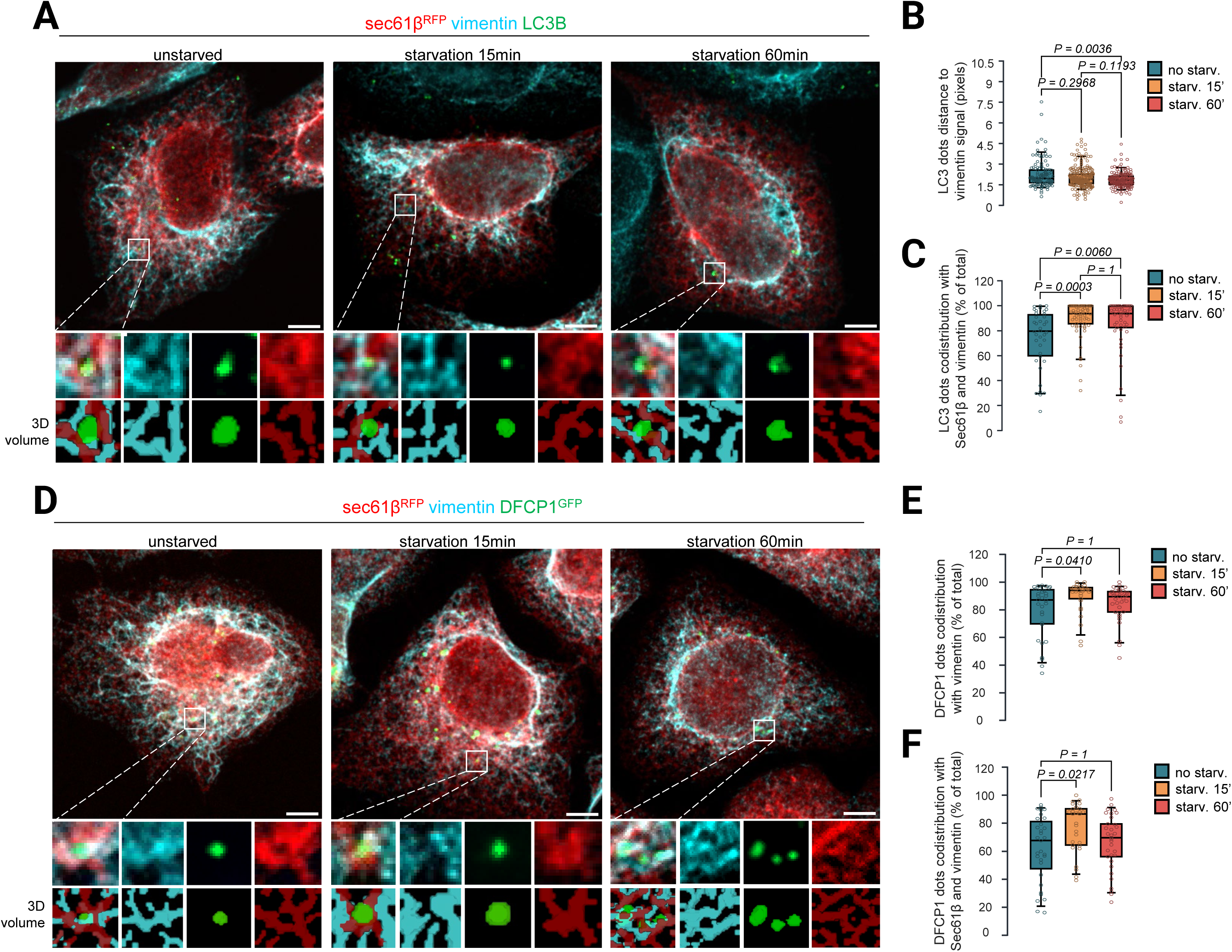
LC3B and DFCP1 are found in proximity to ER and vimentin overlapping regions. HeLa cells were analyzed in basal conditions or after starvation with EBSS for 15 or 60 minutes. (**A**) Representative confocal images of cells transfected with Sec61β-RFP (ER) and labeled with anti-LC3 and anti-vimentin antibodies. Insets show LC3B dots (green) that codistributes with ER (red) and vimentin (cyan) overlapping regions. 3D reconstructions of segmented objects are shown below the raw image. Scale bar = 5 µm. (**B, C**) Quantification of images as shown in (A). (**B**) Number of pixels between LC3B spots and vimentin. N=3 independent experiments; n=132 (basal), 285 (15 min), 195 (60 min) cells. (**C**) Proportion of LC3B spots overlapping with Sec61β-RFP (ER) and vimentin overlapping regions. N=3 independent experiments; n=38 (basal), 65 (15 min), 50 (60 min) cells. For (B) and (C), *P*-values calculated with Kruskal-Wallis test with Dunn’s multiple comparisons test. (**D**) Representative confocal images of cells transfected with Sec61β-RFP (ER) and DFCP1-GFP and labeled with anti-vimentin antibody. Insets show DFCP1-GFP dots (green) that codistributes with ER (red) and vimentin (cyan) overlapping regions. 3D reconstructions of segmented objects are shown below the raw image. Scale bar = 5 µm. (**E, F**) Quantification of images as shown in (D). (**E**) Proportion of DFCP1-GFP overlapping with vimentin. (**F**) Proportion of DFCP1-GFP overlapping with Sec61β-RFP (ER) and vimentin overlapping regions. For (E) and (F), N=3 independent experiments; n=31 (basal), 29 (15 min), 33 (60 min) cells. *P*-values calculated with Kruskal-Wallis test with Dunn’s multiple comparisons test.

### Autophagosome biogenesis events are associated with vimentin structures

Since we detected local vimentin rearrangement associated with ER membrane (Fig.S1) in response to starvation and because ER membrane subdomains serve as active matrix for autophagosome biogenesis (Hu and Reggiori, 2022; Ktistakis, 2020), we analyzed the putative presence of vimentin cytoskeleton at ER-associated pre-autophagic platforms. We performed immunofluorescence to detect LC3B (Fig. 2A), since this protein is commonly used as a *bona-fide* marker to detect autophagic structures (Klionsky et al., 2016). By measuring the distance between LC3-positive puncta and the nearest vimentin object, we observed that LC3 dots were significantly closer to vimentin IFs structures in cells starved for 15 or 60 min. compared to unstarved cells (Fig. 2A, B). Using whole Sec61β^RFP^ (ER) and vimentin segmentation, we observed intricate 3D structures in which LC3 positive vesicles were closely surrounded by ER/vimentin structures (Fig. 2A, 3D reconstructions). We also found that LC3 puncta were significantly closer to the ER-vimentin overlapping regions in cells starved for 60 min. (Fig. 2A, C). LC3B is present not only on phagophore, but also on mature autophagosome and autophagolysosome. To precisely investigate the vimentin relationship with early events of autophagosome biogenesis, we labeled DFCP1, a component selectively associated with the omegasome (Axe et al., 2008). Similar to LC3 signal codistribution with vimentin, we revealed DFCP1 dots inside vimentin and Sec61β^RFP^ (ER) positive meshwork (Fig. 2D, 3D reconstructions). Moreover, the number of these DFCP1/ER/vimentin structures significantly increased in response to starvation, as illustrated by DFCP1 puncta codistribution with vimentin (Fig. 2D, E) and with vimentin and Sec61β^RFP^ overlapping signal (Fig. 2F). We observed a similar pattern for ATG13, a member of the ULK1 complex (Fig. S3), known to initiate autophagosome biogenesis sequence at the ER membrane (Ganley et al., 2009), thus confirming the presence of pre-autophagic proteins in ER/vimentin areas of the cell during response to starvation. These results suggest that autophagosome biogenesis initiation processes, known to take place at ER/omegasome membrane, occur more specifically at areas spatially confined by vimentin IFs.

### Vimentin network is required for starvation-induced autophagy

Given that autophagic and pre-autophagic structures distributed at immediate vicinity of ER/vimentin scaffolds in response to starvation (Fig. 2 and Fig. S3), we thus investigated whether vimentin IF could contribute to autophagic processes in nutrient deprivation. We first analyzed the autophagic flux in starved cells using (or not) withaferin A (WF-A), a drug resulting in a vimentin network destabilization by promoting vimentin aggregation into a dense perinuclear condensate (Bargagna-Mohan et al., 2007). To dynamically assess autophagic flux, we monitored the conversion of LC3B-I to LC3B-II by Western blot in starved HeLa cells, treated with or without 5µM WF-A, in presence or absence of Bafilomycin A1 (BafA1, 100nM), a V-ATPase inhibitor commonly used to block lysosomal degradation to allow the accumulation of lipidated LC3B-II (Klionsky et al., 2016). After 6 hours of starvation, the autophagic flux was significantly lower in WF-A-treated cells compared to the control (Fig. S4A, B). The effect was specific to later time points, as autophagic flux was comparable between control and WF-A-treated cells after 1 hour of starvation and in basal conditions (Fig. 4B). We also analyzed the levels of the autophagy receptor SQSTM1/p62 as a readout of autophagic degradation (Pankiv et al., 2007). Consistently with LC3 data, while p62 levels decreased proportionally to the duration of starvation in control cells, they remained stable in WF-A-treated cells (Fig. S4A, C).

Phosphorylation of ATG16L1 at serine 278 (P-ATG16L1^278S^) was recently described as a highly reliable marker for autophagy induction (Tian et al., 2020). We therefore measured the levels of P-ATG16L1^278S^ in cells treated with 5µM WF-A and starved for 60 or 360 min. P-ATG16L1^278S^ levels accumulated proportionally with the duration of starvation in control cells. In contrast, P-ATG16L1^278S^ levels were significantly decreased in starved cells pre-treated with WF-A compared to control cells (Fig. 3A, B). To go further and avoid any side effects of WF-A treatment, we generated a HeLa cell line knocked-out for vimentin (Fig. 3C, see methods for details). In vimentin KO cells (*VIM*^-/-^), we similarly observed that levels of P-ATG16L1^278S^ were significantly reduced in starvation, compared to wild type cells (Fig. 3D, E). We also found that the formation of LC3 puncta in response to starvation was significantly reduced in vimentin KO cells compared to wild type cells (Fig. S4D, E). Collectively, these results suggest that vimentin contributes to autophagic flux through modulation of early steps of autophagosome biogenesis.

**Figure 3.**
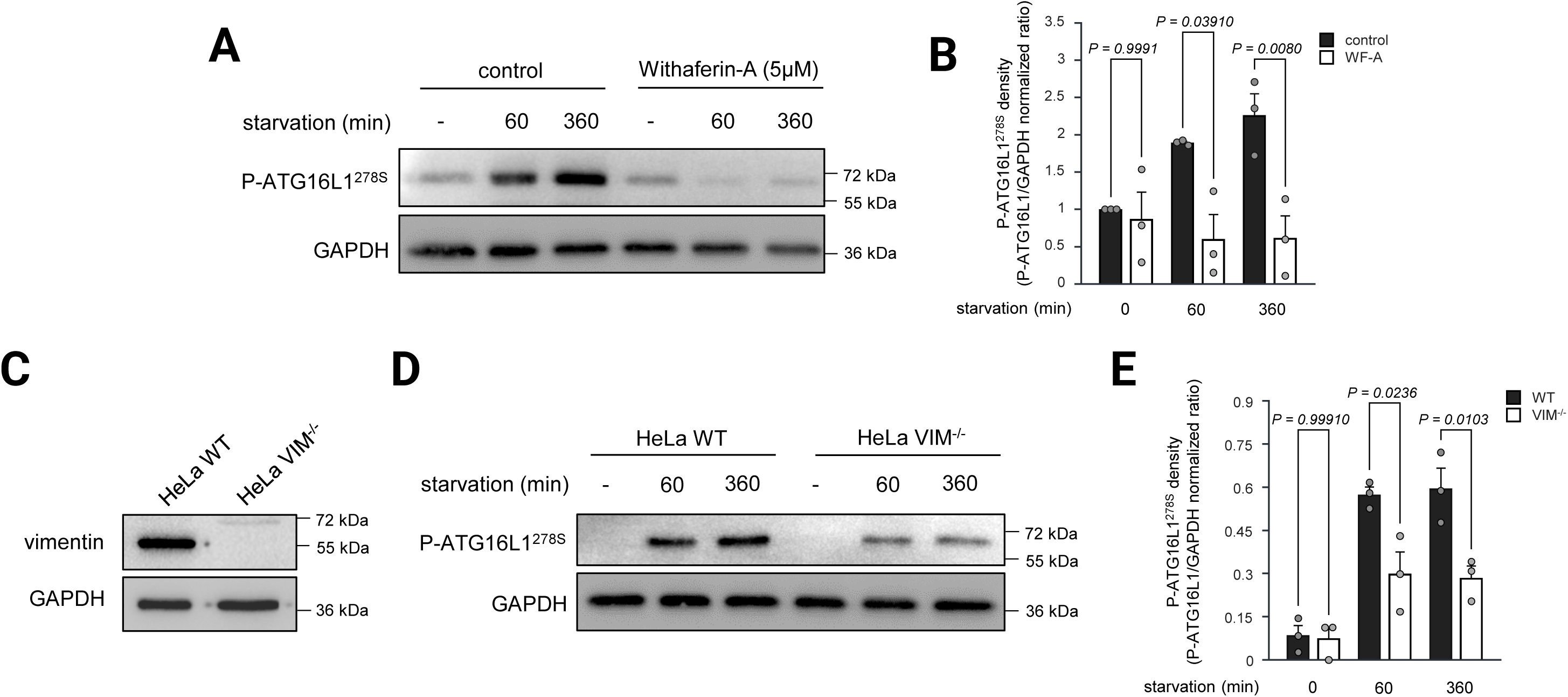
Vimentin destabilization impairs ATG16L1 phosphorylation in response to starvation. (**A, B**) Analysis of pharmacological vimentin destabilization on P-ATG16L1^278S^ in response to starvation. (A) Representative western blot analysis of P-ATG16L1^278S^ and GAPDH expression of HeLa cells treated with or without Withaferin-A (WF-A, 5µM) in basal or EBSS-starved (60 or 360 minutes) conditions. (B) Quantification of western blot analysis shown in (A). P-ATG16L1^278S^ density was normalized with GAPDH density. N=3 independent experiments. *P-*values calculated with Two-way ANOVA with Tukey multiple comparisons test. (**C**) Representative western blot analysis validating vimentin and GAPDH expression in HeLa wild type (WT) or HeLa *VIM^-/-^* cell lines. (**D, E**) Analysis of genetic vimentin knockout destabilization on P-ATG16L1^278S^ in response to starvation. (D) Representative western blot analysis of P-ATG16L1^278S^ and GAPDH expression of HeLa WT or *VIM^-/-^* cells in basal or EBSS-starved (60 or 360 minutes) conditions. (E) Quantification of western blot analysis shown in (D). P-ATG16L1^278S^ density was normalized with GAPDH density. N=3 independent experiments. *P-*values calculated with Two-way ANOVA with Tukey multiple comparisons test.

**Figure 4.**
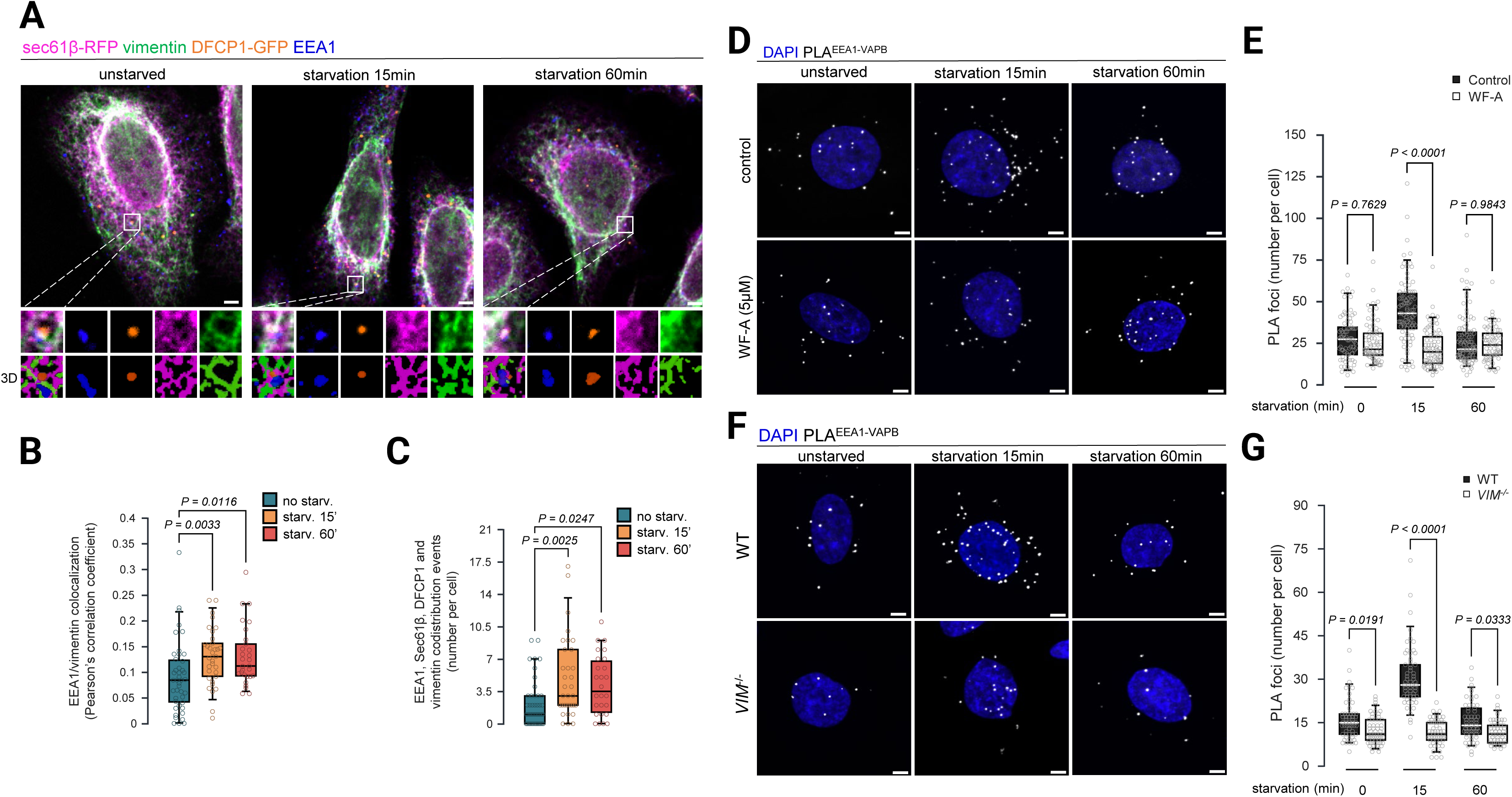
Vimentin promotes ER-early endosomes membrane contact sites in response to starvation. (**A-C**) Vimentin co-distribution with ER, early endosomes, and DFCP1. (**A**) Representative confocal images of HeLa cells transfected with Sec61β-RFP (ER) and DFCP1-GFP and labeled with anti-EEA1 (early endosomes) and anti-vimentin antibodies. Cells were analyzed in basal conditions or after starvation with EBSS for 15 or 60 minutes. Insets show DFCP1-GFP dots (orange) that codistributes with endosomes (blue), ER (magenta) and vimentin (green) overlapping regions. 3D reconstructions of segmented objects are shown below the raw image. Scale bar = 5 µm. (**B, C**) Quantification of images as shown in (**A**). (**B**) Pearson’s correlation coefficient between EEA1 and vimentin signals. N=3 independent experiments; n=43 (basal), 33 (15 min), 29 (60 min) cells. (**C**) Quantification of the number of codistribution events per cell. N=3 independent experiments; n=44 (basal), 33 (15 min), 26 (60 min) cells. For (**B**) and (**C**), P-values calculated with Kruskal-Wallis test with Dunn’s multiple comparisons test. (**D, E**) Pharmacological destabilization of vimentin reduces ER-endosome contacts. (**D**) Representative maximum z projection of confocal images from proximity ligation assays (PLA) using anti-EEA1 and anti-VAPB antibodies in HeLa cells treated with or without WF-A (5µM) in basal condition or after starvation with EBSS for 15 or 60 minutes. Scale bar = 5µm. (**E**) Quantification of number of PLA foci (ER-early endosome membrane contact sites) per cell from (**D**). N=3 independent experiments; n= 78 (control, basal), 93 (control, 15 min), 106 (control, 60 min), 80 (WF-A, basal), 84 (WF-A, 15 min), 72 (WF-A, 60 min). P-values calculated with Kruskal-Wallis test with Dunn’s multiple comparisons test. (**F, G**) Genetic knockout of vimentin reduces ER-endosome membrane contact sites. (**F**) Representative maximum z projection of confocal images from PLA using anti-EEA1 and anti-VAPB antibodies in HeLa WT or VIM-/- cells in basal condition or after starvation with EBSS for 15 or 60 minutes. Scale bar = 5µm. (**G**) Quantification of number of PLA foci per cell from (**F**). **N=3 independent experiments; n= 55 (WT, basal), 57 (WT, 15 min), 57 (WT, 60 min), 61 (VIM-/-, basal), 60 (VIM-/-, 15 min), 56 (VIM-/-, 60 min)**). P-values calculated with Kruskal-Wallis test with Dunn’s multiple comparisons test.

### Vimentin network supports ER-endosomes membrane contact sites in response to starvation

We recently demonstrated that ER–endosome contact sites (EERCS) serve as active platforms for autophagosome biogenesis, a process that requires stabilization of contact tethering and cytoplasmic confinement to enable phagophore initiation (Da Graça et al., 2023; Da Graça et al., 2025). Given the strong accumulation of pre-autophagic markers in regions of close apposition between vimentin and the ER in rapid response to starvation (Fig. 2), we reasoned that vimentin dynamics may contribute to the regulation of EERCS for autophagosome formation.

We first analyzed the codistribution of vimentin with early endosomes (labeled by EEA1) and monitored the formation of ER (Sec61β)–pre-autophagic (DFCP1)–endosome (EEA1)–vimentin structures during starvation (Fig. 4A). Notably, the codistribution between EEA1 and vimentin was significantly increased after 15 min and 60 min of starvation compared to control cells (Fig. 4A, B). Although the absolute correlation values remain low (< 0.2), attributable to the geometric disparity between the extensive filamentous vimentin network and discrete endosomal vesicles, this significant quantitative increase confirms a specific mobilization of endosomes toward the vimentin scaffold during starvation. Importantly, the number of events displaying colocalization of all four markers (EEA1, DFCP1, Sec61β, and vimentin) was also significantly higher in starved cells (Fig. 4A, C). This increase was most pronounced after 15 min of starvation, paralleling the reorganization of the vimentin network induced by nutrient deprivation (Fig. 1 and Fig. S1). Together, these results indicate that local rearrangements of ER–vimentin structures occurring during early autophagosome biogenesis are, at least in part, associated with early endosomes, highlighting a functional relationship between vimentin IF mobilization and EERCS dynamics.

To further investigate EERCS dynamics under these conditions, we performed proximity ligation assays (PLA) with sub-40 nm resolution using antibodies against EEA1 (early endosomes) and VAP-B (ER contact site marker; (Da Graça et al., 2025)) in starved cells, treated or not with 5 µM WF-A (Fig. 4D). While WF-A treatment had no effect on the number of EEA1/VAP-B PLA puncta (i.e., EERCS events) under nutrient-rich conditions, it completely abolished the starvation-induced increase in EERCS observed at early time points (15 min, but not 60 min, as previously reported in (Da Graça et al., 2025)) (Fig. 4D, E). A similar loss of starvation-dependent EERCS dynamics was observed in VIM⁻/⁻ cells (Fig. 4F, G), confirming that vimentin network dynamics are required for starvation-induced stabilization of ER–endosome contacts and supporting a central role for vimentin IFs in ER–endosome communication during autophagosome formation.

## Discussion

The efficient initiation of autophagosome biogenesis requires precise spatial and temporal organization of membranes, yet the cytoskeletal components that orchestrate this dynamic architecture remain incompletely defined (Nambiar and Manjithaya, 2024). Here, we identify the vimentin IF network as a critical cytoskeletal coordinator required for the efficient initiation of starvation-induced autophagic processes. We show that nutrient deprivation triggers a rapid compaction of the vimentin IFs network in the perinuclear area that becomes increasingly associated with the ER. This spatial shift correlates with a transient, localized phosphorylation of vimentin at Serine 56, a modification known to regulate vimentin dynamics (Li et al., 2006). This reorganized vimentin-ER scaffold is associated with a cytoplasmic microenvironment enriched for key autophagic proteins, including DFCP1, ATG13, and LC3B, essential for *de novo* autophagosome biogenesis. We recently established that starvation-associated EERCS are critical hubs for phagophore nucleation, promoting a local environment favoring autophagosome biogenesis (Da Graça et al., 2025). Our present work provides a new layer to that model, demonstrating that the starvation-induced EERCS stabilization is dependent on an intact vimentin network. Functionally, cells lacking vimentin integrity show a significant defect in autophagic flux and, more specifically, a failure to initiate autophagosome biogenesis, as evidenced by impaired ATG16L1 phosphorylation.

The role of the cytoskeleton in autophagy has been intensely studied and the roles of microtubules in facilitating autophagosome-lysosome transport and F-actin in the early stages of phagophore formation are well-established (Kast and Dominguez, 2017; Monastyrska et al., 2009), but the contribution of IFs has remained under investigated, especially in the complex and rapid autophagosome biogenesis sequence. Our data positions vimentin IFs as a distinct, non-redundant player. Thus, beyond the simple mobilization of organelles, we propose that vimentin drives a global re-compartmentalization of the intracellular space, acting not as a transport motor, but as a mechanism of spatial confinement that physically enforces the organelle proximity required for contact site stabilization. We reveal that vimentin IFs do act as a dynamic cytoskeletal coordinator that assembles the membrane platform required for autophagosome biogenesis. This aligns with the emerging view of vimentin as a key mechanical integrator of cellular space and organelle function (Guo et al., 2025; Pradeau-Phélut and Etienne-Manneville, 2024). We show that the tight association of vimentin and the ER, observed decades ago and more recently shown to be critical for ER spreading and reorganization, is harnessed by the cell during starvation to build EERCS (Cremer et al., 2023; Franke et al., 1987; Katsumoto et al., 1990).

Furthermore, the role of vimentin as a "cytoskeletal coordinator" may extend into the biophysical regulation of autophagosome initiation. The assembly of the ULK complex and subsequent recruitment of the PI3KC3-C1 complex are known to be driven by liquid-liquid phase separation (LLPS, (Fujioka et al., 2020; Zheng et al., 2022)). The dense, perinuclear vimentin network and the confined EERCS it organizes appear as ideal microenvironments for promoting such local biophysical processes. By increasing local confinement, the vimentin-EERCS-ER nexus could effectively lower the concentration threshold required for the phase separation of key initiation factors, such as FIP200, ATG13, and ULK1, thereby ensuring that autophagosome biogenesis is nucleated efficiently at the correct membrane site.

The literature on vimentin’s role in autophagy has been, until now, complex and somewhat contradictory. Several studies have proposed an inhibitory role for vimentin, suggesting it sequesters Beclin-1 via a 14-3-3 complex, thereby repressing basal autophagy (Biskou et al., 2019). Conversely, other work has suggested that vimentin is a regulator of autophagy and lysosome positioning, by modulating mTORC1 signaling (Biskou et al., 2019; Mohanasundaram et al., 2022). Vimentin can also be a specific cargo for selective autophagy (Wang et al., 2025). Here, we propose that vimentin IFs integrity is essential for the acute, stress-induced scaffolding of the autophagosome biogenesis machinery.

While our work focuses on vimentin, it raises the possibility that IFs are key spatial regulators of the endolysosomal and autophagic pathways under stress. Indeed, vimentin is known to form cages around organelles like lipid droplets and, through linkers like RNF26, to integrate ER and endolysosomal responses to stress (Franke et al., 1987; Sjödin and Gylfe, 1994). It is likely that other IFs play analogous roles. Interestingly, disruption of the cytokeratin cytoskeleton in hepatocytes or depletion of the type III IF peripherin has also been shown to inhibit autophagic flux (Romano et al., 2025; Seglen et al., 1996; Son et al., 2022), suggesting that IF networks are tailored to spatially organize membranes related to stress, including the autophagic machinery and organelles, in specific cell types and stress contexts.

Although our study establishes a novel vimentin-EERCS axis, it also highlights areas of future investigation. It is in theory possible that the phenotype observed in vimentin null cells arises from a defect in autophagosome-lysosome fusion. Indeed, a defect in lysosomal function is a plausible alternative, particularly given vimentin’s roles in organelle positioning, which could secondarily affect lysosomal-ER contacts for lysosomal homeostasis (Cross et al., 2023; Durgan and Florey, 2022; Romano et al., 2025; Yang et al., 2025). While we cannot completely exclude a secondary role for vimentin in later steps or in lysosome homeostasis, our finding that P-ATG16L1, one of the earliest markers of the initiation cascade, is strongly impaired provides a compelling argument that the first defect is at the initiation steps. Furthermore, this work opens several new mechanistic questions and the precise signaling pathway controlling vimentin compaction remains to be dissected. Moreover, while our data clearly establishes vimentin as an essential coordinator, the molecular machinery that physically tethers the filaments to EERCS components must be identified. Finally, it remains unknown whether vimentin achieves this by acting as a passive scaffold or by actively generating local mechanical force, perhaps in concert with motor proteins.

Altogether, our study reveals that the vimentin IF network is a dynamic regulator of organelle contact sites, by identifying a novel vimentin-EERCS axis that is essential for the rapid initiation of autophagy in response to nutrient stress.

## Material and methods

### Antibodies used in this study

All primary and secondary antibodies used in this study, including host species, sources, catalog numbers, specific dilutions, and Research Resource Identifiers (RRID) are listed in Table S1.

### Cell culture, transfection and treatments

HeLa cells from ATCC were cultured in MEM buffer containing GlutaMAX (Gibco™ 41090036) supplemented with 1% (v/v) non-essential amino acids (Gibco™ 11140050) and 10% (v/v) heat decomplemented fetal calf serum (FCS). Cells were incubated at 37°C, 5% CO2. Cells have been negatively tested for mycoplasma contamination.

Transfections of cDNA plasmids were performed at least 24h before the experiments using FuGENE (E2312; Promega) according to manufacturer’s recommendations. DFCP1-GFP and Tol2 mammalian expression plasmids were from Addgene (#38269 and #31823). Vimentin-eGFP mammalian expression and CRISPR KO plasmids were purchased from Vectorbuilder (VB250304-1119jfg and VB900061-4328cne). Sec61β-RFP cDNA was from T. Rapoport (Boston).

For starvation experiments, cells were washed twice using PBS (Gibco™ 11140050) and incubated in Earle’s Balanced Salt Solution (Gibco™ 24010043) for the indicated times.

### CRISPR/Cas9-mediated knock-out of vimentin in HeLa cells

The guide RNA sequence for CRISPR/Cas9 mediated knockout of human vimentin was designed using CHOPCHOP (https://chopchop.cbu.uib.no/). Sequences targeting VIM: 5’-TTGCTGACGTACGTCACGCA – 3’ were purchased from vectorbuilder.com (VB900061-4328cne) subcloned into pRP-backbone allowing the concomitant expression of hCas9 and the puromycin resistance gene together with eGFP for selection. VB UltraStable (VectorBuilder, UC001-010) bacteria containing the plasmids were grown overnight on agar plates and plasmids were purified from individual colonies (Macherey-Nagel, 740588.250). The purified plasmid was then transfected in HeLa cells using FuGENE (E2312; Promega) according to manufacturer’s recommendations. 24 hours after transfection, cells were treated with puromycin (10µg/mL, Gibco™ A1113803) overnight. Then, surviving GFP positive clones were sorted by fluorescence activated cell sorting (BD FACSAria™ II) in 96 multi well plate. Selected clones were then analyzed for vimentin expression using western blot.

### Western blot analysis

Plated cells were washed 2 times with PBS and lysed with Laemmli buffer (60 mM Tris–HCl pH = 6.8, 2% SDS, 10% glycerol, bromophenol blue) supplemented with 100mM DTT and samples were heated at 95°C for 5 minutes. Protein samples were separated by SDS-PAGE using 13.5% gels in Tris-Glycine-SDS running buffer (Euromedex, EU0510). Following electrophoresis, proteins were transferred to 0.22 µm PVDF membranes (Bio-Rad, 162-0784) using a liquid transfer system. The transfer was performed in Tris-Glycine transfer buffer (Euromedex, EU0550) supplemented with 20% (v/v) absolute ethanol at a constant voltage of 100V for 120 minutes. Blocking was then performed using 5% (m/v) non-fat dry milk diluted in wash buffer (PBST, TBS-0.1% (v/v) Tween 20 (Sigma-Aldrich®, P1379)) for 1 hour at RT. Primary antibodies diluted in PBST-5% (v/m) BSA (Euromedex, 04-100-812) were then incubated overnight at 4°C with shaking. Membranes were washed 3 times for 5 minutes using PBST and secondary antibodies diluted in PBST-5% (v/m) Milk were incubated for 1 hour at RT. Membranes were washed 3 times for 5 minutes using PBST and bands visualized using chemiluminescent HRP substrate (WBKLS0500; MERCK) and a ChemiDoc MP Imaging System (Bio-Rad). Quantification of band intensities was carried out using FiJi.

### Immunofluorescence processing and spinning disk confocal microscopy

For classical immunofluorescence, HeLa cells were seeded on 13mm glass coverslips (No. 1.5H, Deckgläser 0117530). Cells were washed twice with PBS and fixed with 4% (v/v) paraformaldehyde (PFA) diluted in PBS prewarmed to 37°C for 15 minutes. After fixation, cells were washed three times with PBS and blocked with 10% (v/v) FCS in PBS for 30 minutes. Primary antibodies, diluted in staining buffer (PBS supplemented with 10% (v/v) FCS and 0.05% (m/v) saponin), were incubated for 1 hour at room temperature in a humid chamber. Following three 5-minute washes with PBS, secondary antibodies, diluted in staining buffer, were incubated for 1 hour at room temperature. Coverslips were then washed three times for 5 minutes with PBS and mounted with Mowiol, with or without DAPI. After 24h of curing, slides were imaged through a 63x objective (Zeiss 420782-9000, Alpha pln apo DIC, NA=1.4) mounted on a Axio Observer Z1 microscope equipped with a Yokogawa CSU-X1 spinning head and a Prime BSI camera (Teledyne imaging).

### Stimulation emission depletion (STED) microscopy

For STED, HeLa cells were seeded on 13mm glass coverslips (No. 1.5H, Deckgläser 0117530). Cells were then washed twice with PHEM buffer (Electron Microscopy Sciences, 11162) and fixed with 4% (v/v) PFA, 0.2% (v/v) glutaraldehyde (Electron Microscopy Sciences, 16220) and 0.5% (v/v) Triton-X100 diluted in PHEM for 12 minutes at room temperature. After fixation, cells were washed three times with PHEM and blocked with 10% (v/v) FCS in PHEM for 30 minutes. The primary chicken IgY anti-vimentin antibody (1:200, abcam ab24525), diluted in staining buffer (PHEM supplemented with 10% (v/v) FCS and 0.05% (m/v) saponin), was incubated for 1 hour at room temperature in a humid chamber. Following three 5-minute washes with PHEM, the secondary anti-chicken IgY-AF555 (1:200, Invitrogen A21437), diluted in staining buffer, was incubated for 1 hour at room temperature. Coverslips were then washed three times for 5 minutes with PHEM and mounted with Mowiol with DAPI. After 24h of curing, slides were imaged through a 100x objective (HC PL APO 100x/1.40 NA OIL STED WHITE) mounted on a Leica SP8 gSTED system with a white excitation laser, 660nm STED depletion line and a HyD detector. Images were acquired at a 600hz frequency, STED power set to 50% and 3x line averaging with a pixel size of 25nm. Images were then further deconvolved with the deconvolution express wizard using standard settings within Huygens Professional 25.10 (Scientific Volume Imaging).

### Proximity ligation assay (PLA)

For proximity ligation assays (PLA), cells were seeded on 13mm glass coverslips and fixed for 15 minutes at 37°C with 4% (v/v) PFA diluted in PBS. After three washes with PBS, cells were permeabilized with 0.2% (v/v) Triton-X100 in PBS for 10 minutes at room temperature. All PLA experiments were performed using Duolink PLA kits (Sigma-Aldrich, DUO92101) according to the manufacturer’s instructions. Following three washes with PBS, samples were blocked for 1 hour at 37°C in a humid chamber with Duolink blocking solution. Primary antibodies were diluted in Duolink antibody diluent and incubated with the cells for 1 hour at room temperature. After three washes with PBS, cells were incubated with the corresponding Duolink secondary antibodies conjugated with a PLUS and MINUS end for 1 hour at room temperature. The ligase reaction was then performed by incubating the cells for 30 minutes at 37°C with Duolink ligase diluted in Duolink Ligation Buffer. After three washes with PBS, cells were incubated for 100 minutes at 37°C with Duolink polymerase diluted in Duolink Polymerase Buffer. Finally, following three washes with PBS, the coverslips were mounted with Mowiol and allowed to be cured for 24 hours at RT before imaging.

### Image analysis

Image analyses were conducted using FiJi, CellProfiler 4.2.8 or napari. Object segmentations were performed on each separated channel using Otsu three class method in the IdentifyPrimaryObjects CellProfiler module or the Nellie plugin for Napari. Cell segmentation was performed by the IdentifySecondaryObjects module using the raw vimentin channel and the propagation method using segmented nuclei as centers. Following segmentation, new sets of objects were created by RelateObjects or MaskObjects modules. Objects morphological features (Area, compactness indices) of each object were measured by the MeasureSizeAndShape module. The fraction of perinuclear vimentin was calculated by the MeasureIntensityDistribution module within each segmented cell and using the edge of segmented nuclei. Similarly, colocalization coefficients (Pearson’s, Mander’s) were calculated using raw channels within each segmented cell with the MesureColocalization module. 3D reconstructions were conducted by overlaying segmented objects in napari.

### Statistical analysis

Normality of datasets were tested using the Shapiro-Wilk test of normality. Accordingly, non-parametric and statistical analysis were performed using Kruskal-Wallis with Dunn’s multiple comparisons test, two-way ANOVA with Tukey multiple comparisons test or Mann-Whitney test using the rstatix package running on R 4.2.2. All statistical analyses were performed from at least three independent experiments and statistical significance was considered for p values <0.05.

## Acknowledgements and funding

QF performed all the biochemical and imaging experiments, formal analysis, quantifications, data analyses and wrote the manuscript. DL contributed to some of the biochemical experiments. CD contributed to supervision, formal analysis and writing of the manuscript. EM supervised the project and the methodology, contributed to formal analysis and wrote the manuscript.

We are grateful to our colleague Aurore Claude-Taupin from INEM (Paris, France) for critical reading of the manuscript and helpful suggestions. We thank Necker SFR technical platforms and especially Béatrice Durel (imaging platform) for STED microscopy. EM and CD received financial support from INSERM (Institut National pour la Santé et la Recherche Médicale), CNRS (Centre National pour la Recherche Scientifique), Université-Paris Cité, French National Research Agency (grants ANR 22-CE14-0019 and ANR-23-CE14-0041-01), and French Foundation for Medical Research (FRM, « labélisation équipe »).

---

**Figure sup. 1.**
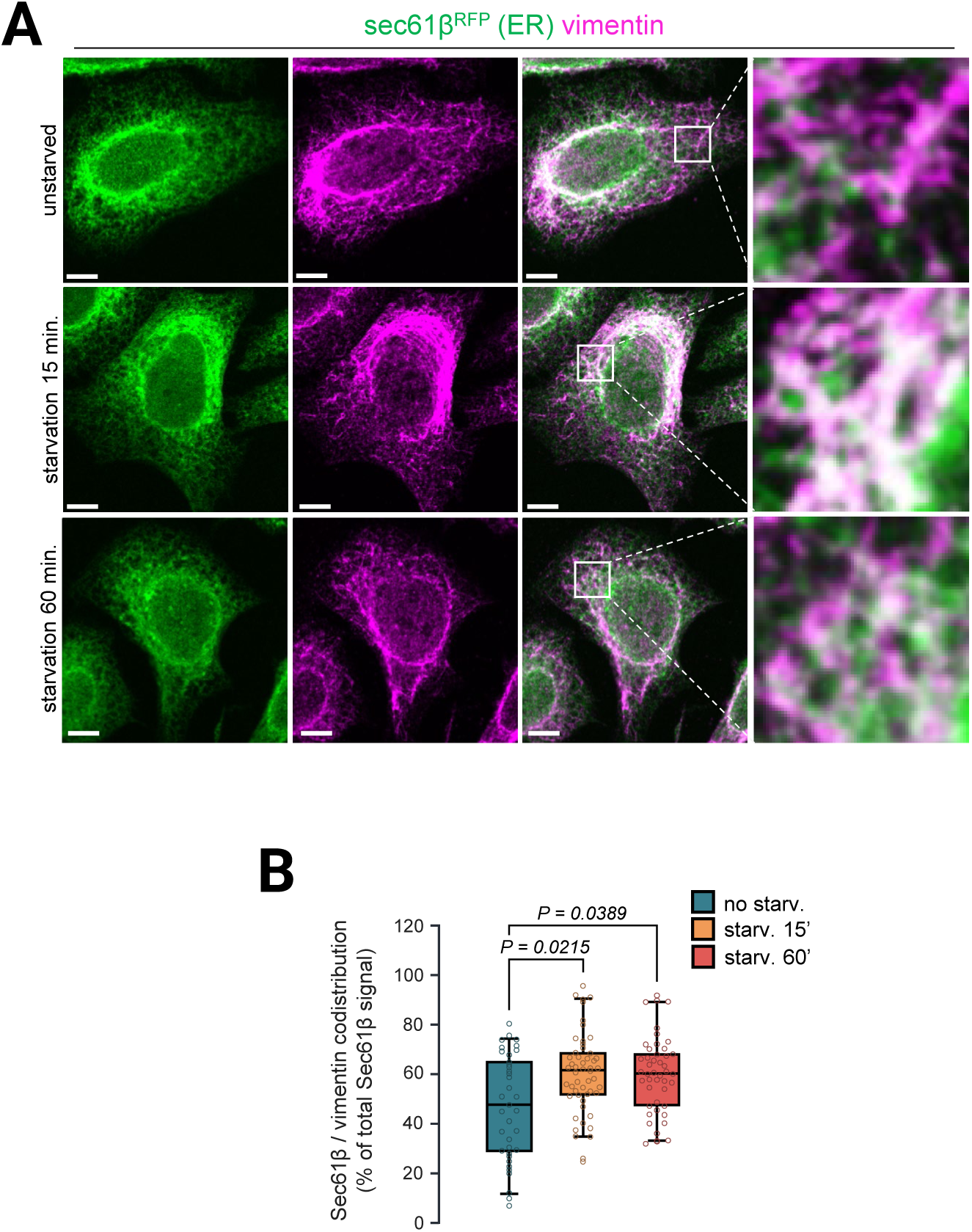
Vimentin overlap with the ER increases in response to starvation. HeLa cells were analyzed in basal conditions or after starvation with EBSS for 15 or 60 minutes. (**A**) Representative confocal images of cells transfected with Sec61β-RFP (ER) and labeled with anti-vimentin antibody. Scale bar = 5 µm. (**B**) Quantification of the proportion of Sec61β-RFP (ER) overlapping with vimentin from images as shown in (**A**). N= 3 independent experiments; n= 37 (basal), 52 (15 min), 46 (60 min) cells. *P*-values calculated with Kruskal-Wallis test with Dunn’s multiple comparisons test.

**Figure sup. 2.**
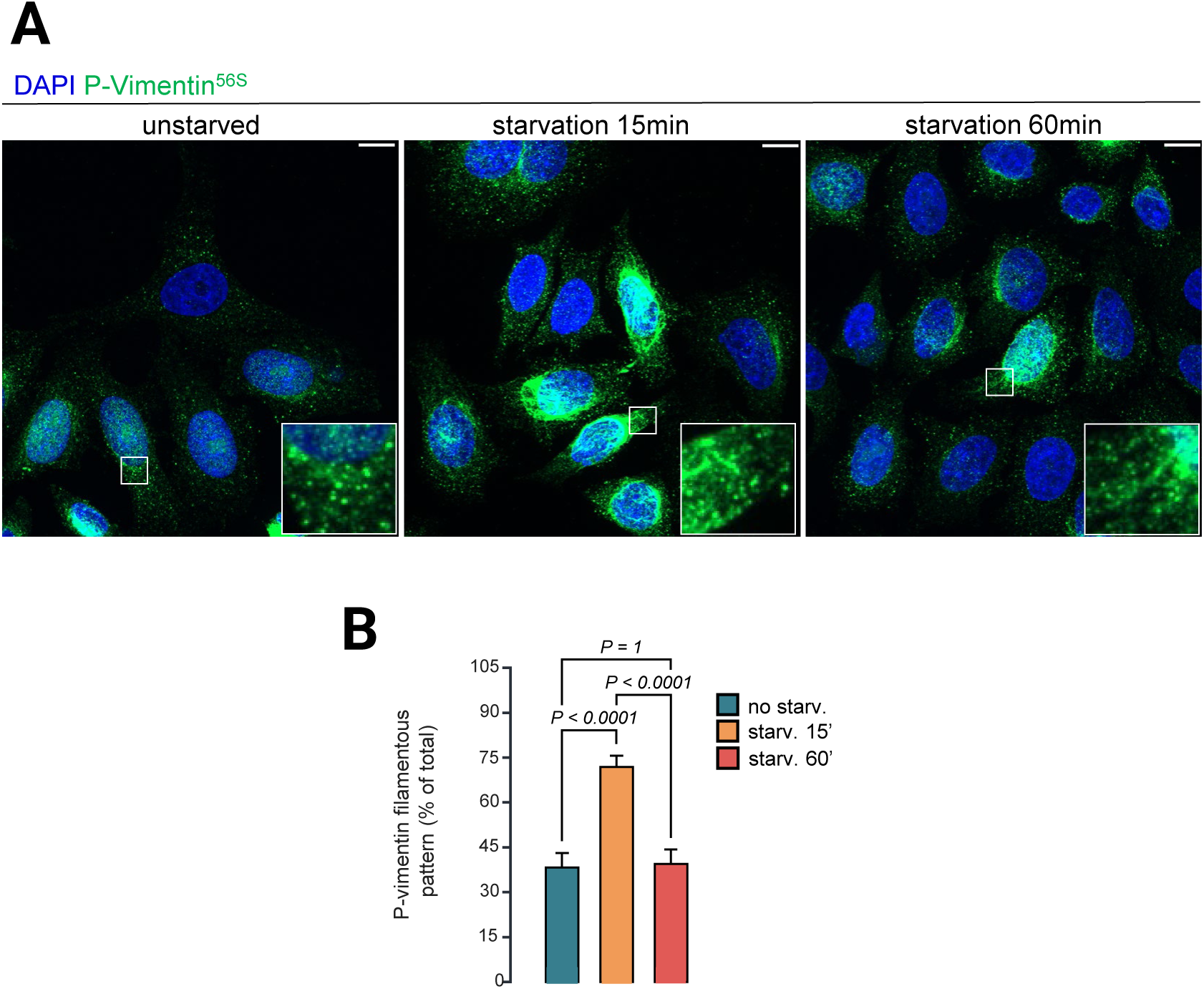
Vimentin phosphorylation at serine 56 is dynamically regulated in response to starvation. HeLa cells were analyzed in basal conditions or after starvation with EBSS for 15 or 60 minutes. (**A**) Representative confocal images of cells labeled with anti-P-Vim56S antibody and DAPI. Insets show “dotty” (basal, 60 min) or “filamentous” patterns (15 min). Scale bar = 5 µm. (**B**) Quantification of the proportion of cells displaying a filamentous P-Vim56S pattern. N= 3 independent experiments; n= 102 (basal), 142 (15 min), 104 (60 min) cells. P-values calculated with Kruskal-Wallis test with Dunn’s multiple comparisons test.

**Figure sup. 3:**
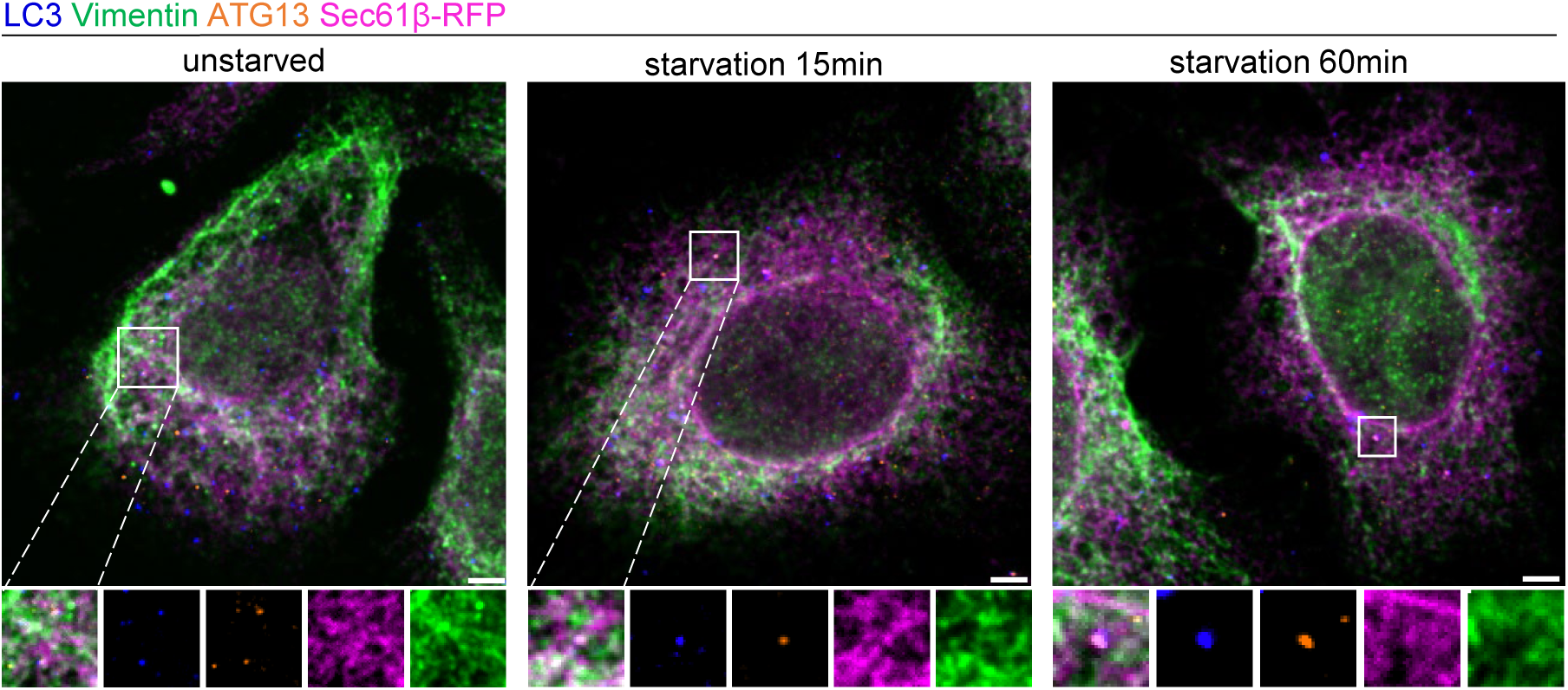
ATG13 co-distributes with LC3 in ER and vimentin overlapping regions in response to starvation. (**A**) Representative confocal images of HeLa cells transfected with Sec61β-RFP (ER) and labeled with anti-vimentin, anti-ATG13 and anti-LC3B antibodies in basal conditions or after starvation with EBSS for 15 or 60 minutes. Scale bar = 5 µm. Insets show ATG13 overlapping with LC3 and codistributing with ER and vimentin overlapping regions.

**Figure sup. 4.**
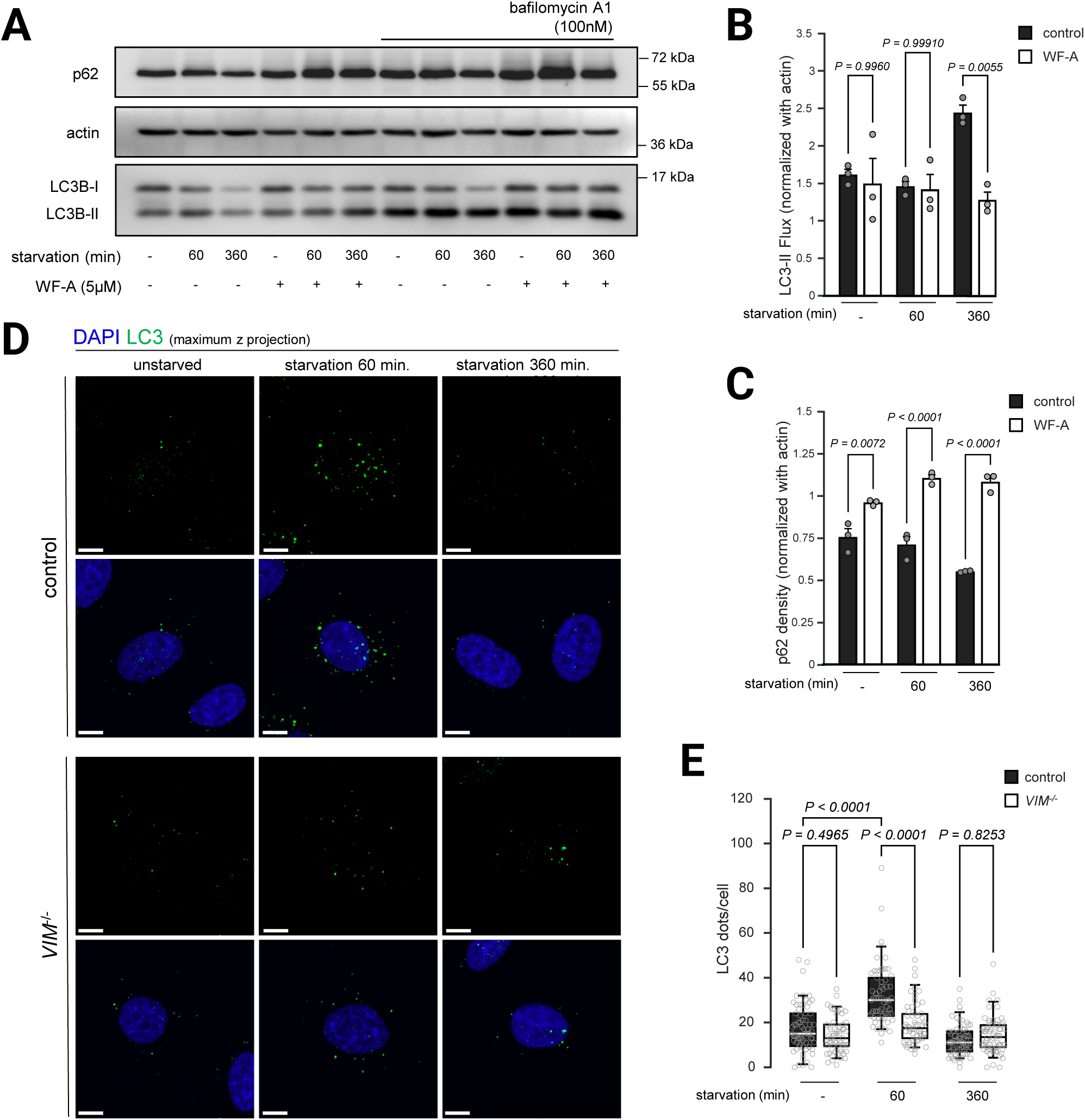
Pharmacological destabilization of vimentin by Withaferin-A treatment impairs autophagic flux. (**A**) Representative western blot analysis of p62, LC3B and actin expression of HeLa cells treated with or without Withaferin-A (WF-A, 5µM) and with or without bafilomycin A1 (100 nM) in basal or EBSS-starved (60 or 360 minutes) conditions. (**B** and **C**) Quantification of western blot analysis shown in (A). (**B**) LC3B-II density was normalized with actin density. For each starvation timepoint, LC3B-II flux was calculated as the LC3B-II density of the condition with bafilomycin A1 normalized with the LC3B-II density of the condition without bafilomycin A1. (**C**) p62 density was normalized with actin density. N=3 independent experiments. *P-*values calculated with Two-way ANOVA with Tukey multiple comparisons test. (**D**) Representative maximum z projection of confocal images of HeLa cells (control and vimentin KO) labeled with anti-LC3 and DAPI in basal of after starvation with EBSS for 60 or 360 minutes. Scale bar = 5µm. (**E**) Quantification of the number of LC3 puncta per cell of images shown in (D). N=3 independent experiments; n= 67 (WT, basal), 61 (WT, 60min), 65 (WT, 360min), 60 (VIM-/-, basal), 58 (VIM-/-, 60min), 66 (VIM-/-, 360min) cells. P-values calculated with Kruskal-Wallis test with Dunn’s multiple comparisons test.

## References

1. Axe, E. L., Walker, S. A., Manifava, M., Chandra, P., Roderick, H. L., Habermann, A., Griffiths, G., & Ktistakis, N. T. (2008). Autophagosome formation from membrane compartments enriched in phosphatidylinositol 3-phosphate and dynamically connected to the endoplasmic reticulum. J Cell Biol, 182(4), 685–701. http://www.ncbi.nlm.nih.gov/entrez/query.fcgi?cmd=Retrieve&db=PubMed&dopt=Citatio n&list_uids=18725538

2. Bargagna-Mohan, P., Hamza, A., Kim, Y. eon, Khuan Abby Ho, Y., Mor-Vaknin, N., Wendschlag, N., Liu, J., Evans, R. M., Markovitz, D. M., Zhan, C. G., Kim, K. B., & Mohan, R. (2007). The Tumor Inhibitor and Antiangiogenic Agent Withaferin A Targets the Intermediate Filament Protein Vimentin. Chemistry and Biology, 14(6), 623–634. 10.1016/j.chembiol.2007.04.010

3. Barlan, K., & Gelfand, V. I. (2017). Microtubule-Based Transport and the Distribution, Tethering, and Organization of Organelles. Cold Spring Harbor Perspectives in Biology, 9(5). 10.1101/CSHPERSPECT.A025817

4. Battaglia, R. A., Delic, S., Herrmann, H., & Snider, N. T. (2018). Vimentin on the move: New developments in cell migration. F1000 Research, 7. 10.12688/F1000RESEARCH.15967.1/DOI

5. Biskou, O., Casanova, V., Hooper, K. M., Kemp, S., Wright, G. P., Satsangi, J., Barlow, P. G., & Stevens, C. (2019). The type III intermediate filament vimentin regulates organelle distribution and modulates autophagy. PLoS ONE, 14(1). 10.1371/journal.pone.0209665

6. Brown, S. S. (1999). Cooperation between microtubule- and actin-based motor proteins. Annual Review of Cell and Developmental Biology, 15, 63–80. 10.1146/ANNUREV.CELLBIO.15.1.63

7. Cremer, T., Voortman, L. M., Bos, E., Jongsma, M. L., Ter Haar, L. R., Akkermans, J. J., Talavera Ormeño, C. M., Wijdeven, R. H., de Vries, J., Kim, R. Q., Janssen, G. M., van Veelen, P. A., Koning, R. I., Neefjes, J., & Berlin, I. (2023). RNF26 binds perinuclear vimentin filaments to integrate ER and endolysosomal responses to proteotoxic stress. The EMBO Journal, 42(18). 10.15252/EMBJ.2022111252

8. Cross, J., Durgan, J., McEwan, D. G., Tayler, M., Ryan, K. M., & Florey, O. (2023). Lysosome damage triggers direct ATG8 conjugation and ATG2 engagement via non-canonical autophagy. The Journal of Cell Biology, 222(12). 10.1083/JCB.202303078

9. Da Graça, J., Charles, J., Djebar, M., Alvarez-Valadez, K., Botti, J., Morel, E. (2023). A SNX1-SNX2-VAPB partnership regulates endosomal membrane rewiring in response to nutritional stress. Life Science Alliance, 6(3). 10.26508/lsa.202201652

10. Da Graça, J., Delevoye, C., & Morel, E. (2024). Morphodynamical adaptation of the endolysosomal system to stress. The FEBS Journal. 10.1111/FEBS.17154

11. Da Graça, J., Thiola, C., Rouabah, M., Guerrera, I. C., El Khallouki, N., Romao, M., Alemany, C., Amiri, D., Dubois, S., Srimoorthy, A., Giordano, F., Raposo, G., & Morel, E. (2025). ER-endosome contacts generate a local environment promoting phagophore formation. Cell Reports, 44(7). 10.1016/j.celrep.2025.115993

12. Dooley, H. C., Razi, M., Polson, H. E. J., Girardin, S. E., Wilson, M. I., & Tooze, S. a. (2014). WIPI2 links LC3 conjugation with PI3P, autophagosome formation, and pathogen clearance by recruiting Atg12-5-16L1. Mol Cell, 55(2), 238–252. 10.1016/j.molcel.2014.05.021

13. Durgan, J., & Florey, O. (2022). Many roads lead to CASM: Diverse stimuli of noncanonical autophagy share a unifying molecular mechanism. Science Advances, 8(43). 10.1126/SCIADV.ABO1274

14. Eriksson, J. E., Dechat, T., Grin, B., Helfand, B., Mendez, M., Pallari, H. M., & Goldman, R. D. (2009). Introducing intermediate filaments: from discovery to disease. The Journal of Clinical Investigation, 119(7), 1763–1771. 10.1172/JCI38339

15. Franke, W. W., Hergt, M., & Grund, C. (1987). Rearrangement of the vimentin cytoskeleton during adipose conversion: Formation of an intermediate filament cage around lipid globules. Cell, 49(1), 131–141. 10.1016/0092-8674(87)90763-X

16. Fujioka, Y., Alam, J. Md., Noshiro, D., Mouri, K., Ando, T., Okada, Y., May, A. I., Knorr, R. L., Suzuki, K., Ohsumi, Y., & Noda, N. N. (2020). Phase separation organizes the site of autophagosome formation. Nature, 578(7794), 301–305. 10.1038/s41586-020-1977-6

17. Ganley, I. G., Lam, D. H., Wang, J., Ding, X., Chen, S., & Jiang, X. (2009). ULK1·ATG13·FIP200 complex mediates mTOR signaling and is essential for autophagy. Journal of Biological Chemistry, 284(18), 12297–12305. 10.1074/jbc.M900573200

18. Guo, M., Wong, I. Y., Moore, A. S., Medalia, O., Lippincott-Schwartz, J., Weitz, D. A., & Goldman, R. D. (2025). Vimentin intermediate filaments as structural and mechanical coordinators of mesenchymal cells. Nature Cell Biology, 27(8), 1210–1218. 10.1038/S41556-025-01713-X

19. Hamasaki, M., Furuta, N., Matsuda, A., Nezu, A., Yamamoto, A., Fujita, N., Oomori, H., Noda, T., Haraguchi, T., Hiraoka, Y., Amano, A., & Yoshimori, T. (2013). Autophagosomes form at ER-mitochondria contact sites. Nature, 495(7441), 389–393. 10.1038/nature11910

20. Hu, Y., & Reggiori, F. (2022). Molecular regulation of autophagosome formation. Biochemical Society Transactions, 50(1), 55–69. 10.1042/BST20210819

21. Kast, D. J., & Dominguez, R. (2017). The Cytoskeleton–Autophagy Connection. Current Biology, 27(8), R318–R326. 10.1016/j.cub.2017.02.061

22. Katsumoto, T., Mitsushima, A., & Kurimura, T. (1990). The role of the vimentin intermediate filaments in rat 3Y1 cells elucidated by immunoelectron microscopy and computer-graphic reconstruction. Biology of the Cell, 68(2), 139–146. 10.1016/0248-4900(90)90299-I

23. Kim, J., Kundu, M., Viollet, B., & Guan, K.-L. (2011). AMPK and mTOR regulate autophagy through direct phosphorylation of Ulk1. Nature Cell Biology, 13(2), 132–141. 10.1038/ncb2152

24. Klionsky, D. J., Abdelmohsen, K., Abe, A., Abedin, M. J., Abeliovich, H., Acevedo Arozena, A., Adachi, H., Adams, C. M., Adams, P. D., Adeli, K., Adhihetty, P. J., Adler, S. G., Agam, G., Agarwal, R., Aghi, M. K., Agnello, M., Agostinis, P., Aguilar, P. V, Aguirre-Ghiso, J., … Zughaier, S. M. (2016). Guidelines for the use and interpretation of assays for monitoring autophagy (3rd edition). Autophagy, 12(1), 1–222. 10.1080/15548627.2015.1100356

25. Klionsky, D. J., Petroni, G., Amaravadi, R. K., Baehrecke, E. H., Ballabio, A., Boya, P., Bravo-San Pedro, J. M., Cadwell, K., Cecconi, F., Choi, A. M. K., Choi, M. E., Chu, C. T., Codogno, P., Colombo, M. I., Cuervo, A. M., Deretic, V., Dikic, I., Elazar, Z., Eskelinen, E., … Pietrocola, F. (2021). Autophagy in major human diseases. The EMBO Journal, 40(19). 10.15252/EMBJ.2021108863

26. Ktistakis, N. T. (2020). ER platforms mediating autophagosome generation. Biochimica et Biophysica Acta (BBA) - Molecular and Cell Biology of Lipids, 1865(1). 10.1016/j.bbalip.2019.03.005

27. Kumar, Y., & Valdivia, R. H. (2008). Actin and Intermediate Filaments Stabilize the Chlamydia trachomatis Vacuole by Forming Dynamic Structural Scaffolds. Cell Host and Microbe, 4(2), 159–169. 10.1016/j.chom.2008.05.018

28. Li, Q. F., Spinelli, A. M., Wang, R., Anfinogenova, Y., Singer, H. A., & Tang, D. D. (2006). Critical role of vimentin phosphorylation at Ser-56 by p21-activated kinase in vimentin cytoskeleton signaling. Journal of Biological Chemistry, 281(45), 34716–34724. 10.1074/jbc.M607715200

29. Mizushima, N., & Levine, B. (2020). Autophagy in Human Diseases. New England Journal of Medicine, 383(16), 1564–1576. 10.1056/nejmra2022774

30. Mohanasundaram, P., Coelho-Rato, L. S., Modi, M. K., Urbanska, M., Lautenschläger, F., Cheng, F., & Eriksson, J. E. (2022). Cytoskeletal vimentin regulates cell size and autophagy through mTORC1 signaling. PLoS Biology, 20(9). 10.1371/JOURNAL.PBIO.3001737

31. Monastyrska, I., Rieter, E., Klionsky, D. J., & Reggiori, F. (2009). Multiple roles of the cytoskeleton in autophagy. Biological Reviews of the Cambridge Philosophical Society, 84(3), 431–448. 10.1111/J.1469-185X.2009.00082.X

32. Nambiar, A., & Manjithaya, R. (2024). Driving autophagy - the role of molecular motors. Journal of Cell Science, 137(3). 10.1242/JCS.260481

33. Nascimbeni, A. C., Giordano, F., Dupont, N., Grasso, D., Vaccaro, M. I., Codogno, P., & Morel, E. (2017). ER-plasma membrane contact sites contribute to autophagosome biogenesis by regulation of local PI3P synthesis. The EMBO Journal, 36(14), e201797006. 10.15252/embj.201797006

34. Pankiv, S., Clausen, T. H., Lamark, T., Brech, A., Bruun, J. A., Outzen, H., Øvervatn, A., Bjørkøy, G., & Johansen, T. (2007). p62/SQSTM1 binds directly to Atg8/LC3 to facilitate degradation of ubiquitinated protein aggregates by autophagy*[S]. Journal of Biological Chemistry, 282(33), 24131–24145. 10.1074/jbc.M702824200

35. Pasolli, M., Meiring, J. C. M., Conboy, J. P., Koenderink, G. H., & Akhmanova, A. (2025). Optogenetic and chemical genetic tools for rapid repositioning of vimentin intermediate filaments. The Journal of Cell Biology, 224(9). 10.1083/JCB.202504004

36. Polson, H. E. J. J., de Lartigue, J., Rigden, D. J., Reedijk, M., Urbé, S., Clague, M. J., Tooze, S. A., Urbe, S., Clague, M. J., & Tooze, S. A. (2010). Mammalian Atg18 (WIPI2) localizes to omegasome-anchored phagophores and positively regulates LC3 lipidation. Autophagy, 6(4), 506–522. 10.4161/auto.6.4.11863

37. Pradeau-Phélut, L., & Etienne-Manneville, S. (2024). Cytoskeletal crosstalk: A focus on intermediate filaments. Current Opinion in Cell Biology, 87. 10.1016/j.ceb.2024.102325

38. Romano, R., Cordella, P., & Bucci, C. (2025). The Type III Intermediate Filament Protein Peripherin Regulates Lysosomal Degradation Activity and Autophagy. International Journal of Molecular Sciences, 26(2). 10.3390/IJMS26020549

39. Russell, R. C., Tian, Y., Yuan, H., Park, H. W., Chang, Y. Y., Kim, J., Kim, H., Neufeld, T. P., Dillin, A., & Guan, K. L. (2013). ULK1 induces autophagy by phosphorylating Beclin-1 and activating VPS34 lipid kinase. Nature Cell Biology, 15(7), 741–750. 10.1038/NCB2757

40. Seglen, P. O., Berg, T. O., Blankson, H., Fengsrud, M., Holen, I., & Strømhaug, P. E. (1996). Structural aspects of autophagy. Advances in Experimental Medicine and Biology, 389, 103–111. http://www.ncbi.nlm.nih.gov/pubmed/8860999

41. Sjödin, L., & Gylfe, E. (1994). Down-regulation of bombesin binding to guinea-pig pancreatic acinar cells during homologous desensitization. British Journal of Pharmacology, 112(4), 1037–1042. 10.1111/j.1476-5381.1994.tb13187.x

42. Son, S., Baek, A., Lee, J. H., & Kim, D. E. (2022). Autophagosome-lysosome fusion is facilitated by plectin-stabilized actin and keratin 8 during macroautophagic process. Cellular and Molecular Life Sciences: CMLS, 79(2). 10.1007/S00018-022-04144-1

43. Teo, C. S. H., & Chu, J. J. H. (2014). Cellular vimentin regulates construction of dengue virus replication complexes through interaction with NS4A protein. Journal of Virology, 88(4), 1897–1913. 10.1128/JVI.01249-13

44. Tian, W., Alsaadi, R., Guo, Z., Kalinina, A., Carrier, M., Tremblay, M. E., Lacoste, B., Lagace, D., & Russell, R. C. (2020). An antibody for analysis of autophagy induction. Nature Methods, 17(2), 232–239. 10.1038/S41592-019-0661-Y

45. Wang, H., Hu, J., Wang, D., Cai, Y., Zhu, W., Deng, R., Zhang, Y., Dong, Z., Yang, Z., Xiao, J., Li, A., & Liu, Z. (2025). TM9SF1 inhibits colorectal cancer metastasis by targeting Vimentin for Tollip-mediated selective autophagic degradation. Cell Death and Differentiation, 32(10). 10.1038/S41418-025-01498-4

46. Yang, H., Xun, J., Li, Y., Mondal, A., Lv, B., Watkins, S. C., Shi, L., & Tan, J. X. (2025). LYVAC/PDZD8 is a lysosomal vacuolator. *Science (New York*, N.Y.), 389(6762). 10.1126/SCIENCE.ADZ0972

47. Young, A. R. J., Chan, E. Y. W., Hu, X. W., Köchl, R., Crawshaw, S. G., High, S., Hailey, D. W., Lippincott-Schwartz, J., & Tooze, S. A. (2006). Starvation and ULK1-dependent cycling of mammalian Atg9 between the TGN and endosomes. Journal of Cell Science, 119(Pt 18), 3888–3900. 10.1242/jcs.03172

48. Zheng, Q., Chen, Y., Chen, D., Zhao, H., Feng, Y., Meng, Q., Zhao, Y., & Zhang, H. (2022). Calcium transients on the ER surface trigger liquid-liquid phase separation of FIP200 to specify autophagosome initiation sites. Cell, 185(22), 4082–4098.e22. 10.1016/j.cell.2022.09.001

